# Long-term co-maturation of stem cell-derived microglia and neuronal networks: an optimized platform to assess human microglial contribution to neuronal function

**DOI:** 10.1101/2025.10.24.684345

**Authors:** Annika Mordelt, Imke M. E. Schuurmans, Nicky Scheefhals, Marina P. Hommersom, Koen Slottje, Kimberly Mast, Mara Graziani, Carlos O. González, Laura J. A. Wingens, Dirk Schubert, Nael Nadif Kasri, Lot D. de Witte

## Abstract

Microglia-neuron interactions play a central role in a variety of central nervous system disorders. Technologies using human induced pluripotent stem cells (hiPSCs) have been developed to model human brain cells with the goal to understand their function. To effectively study neuro-immune crosstalk and investigate microglial contributions to neuronal network development and function, both microglia and neurons should co-mature allowing for long-term interactions throughout their differentiation. Here, we present a co-maturation protocol that robustly generates glutamatergic neuronal networks containing human iPSC-derived microglia. We validated the long-term co-cultures using single-cell transcriptomics, imaging and neuronal activity readouts.

We show that astrocytes were required for long-term survival of microglia and for their integration into neuronal networks. Our co-maturation approach induced the typical ramified microglia morphology and characteristic microglia-neuron interactions. Homeostatic markers like P2RY12 and TMEM119 and neuronal remodeling associated genes were upregulated compared to microglia monocultures, highlighting the necessity of the environment to generate and maintain the context-dependent microglia signature *in vitro.* In this manuscript, we include the full optimization process of our co-maturation approach, a comprehensive description of the protocol, practical guidelines and troubleshooting tips. Our co-maturation model provides a powerful tool to assess the role of human microglia in modulating neuronal function and development in health and disease.

**Highlights:**

- ptimized human iPSC differentiation protocol that allows for co-maturation of microglia and *Neurogenin2*-induced neuronal networks.
- are required for long-term survival and integration of microglia into neuronal networks.
- Co-maturation approach enables characteristic neuron-microglia interactions and induces signature morphology, transcriptome and proteins of human microglia.

## Introduction

Neuro-immune interactions are increasingly recognized to be critical for brain function in health and disease. Microglia, the tissue-resident immune cells of the brain, are at the intersection of neuronal and immunological activity in the central nervous system (CNS)^1,2^. By continuously surveilling the CNS, microglia shape and control neuronal circuits^3–5^. Besides the more classical myeloid functions in response to danger signals, such as initiating inflammatory responses and phagocytosis, the continuous and dynamic interaction with neurons is thought to be central to microglial function and the CNS. Mechanisms like synaptic remodeling have been repeatedly highlighted in the field for the last decade^6–9^, and speculated to be a major mechanism involved in neurodevelopmental, neuroinflammatory and neurodegenerative disorders^10–13^. Therefore, there is an ongoing effort to understand microglial function, microglia-neuron crosstalk and its role in CNS disease.

Unraveling these processes is challenged by the following factors: 1) Microglia adopt diverse states and their function is context dependent^14–17^, we need models that recapitulate the *in vivo* microglia phenotypes. 2) Microglia function and glia-neuron communication is species-specific^16,18^, we need human models to complement rodent *in vivo* models. 3) Microglia-neuron interactions are fast and dynamic, based on their regional context. Models should be able to measure microglia-neuron interactions over time. 4) These interactions take place at the subcellular level. Models should offer the resolution to analyze neuron-microglia interactions at the level of dendrites and synapses. 5) The ultimate output of these interactions is neuronal function. We should build models that can measure the impact of neuron-microglia interactions on neuronal network activity. Current models including microglial cell lines, post-mortem tissue, rodent models, and human induced pluripotent stem cell (hiPSC)-models have been valuable to the field, with their own benefits and challenges. To complement these models we optimized a hiPSC-derived neuron-microglia co-maturation model that addresses all of the five challenges described above.

The discovery of iPSCs^19^ and their use for *in vitro* modeling of CNS disorders yielded opportunities for the neuroimmunology field. Many advances in human microglia modeling have been made^20,21^. Still, difficulties in maintaining the microglia signature *in vitro* remain. Without continuous input of the brain environment, microglia rapidly lose their characteristic phenotype, including a downregulated expression of key markers and a change of their morphology^16,22^. This environment-dependent nature of microglia poses significant challenges for accurate *in vitro* modeling. As a result, monoculture systems, whether based on primary microglia or those derived from hiPSCs, are limited in their ability to replicate the physiological properties of microglia. Further, interactions with neurons are fundamental to microglial physiology and pathology^8,9,12^. Investigating how microglia regulate neuronal circuits requires models that recapitulate the dynamic and continuous interplay between both cell types.

In the last decade, co-cultures with neurons or (xenografted) organoids have been developed^21,23^. However, most current protocols employ separate differentiation of microglia and neurons, followed by a brief co-culture of 24 hours^24,25^ or three days^26^. For comprehensive and robust studies of microglia-neuron interactions including neurodevelopmental aspects, both long-term co-culturing as well as co-maturation of all cell types are required. Microglia-organoid models have been used for such co-maturation^23^ but they are cost- and time-consuming due to long maturation times. Furthermore, the need for xenotransplantation of the organoids to induce the required microglial signature is limiting high-throughput studies, drug screening, and counteracting a reduction of animal use in hiPSC-based research. In contrast, a human-based 2D model provides advantages for studying microglia-neuron crosstalk like high temporal and spatial resolution, possibilities to manipulate and measure at both cellular and subcellular level, and longitudinal non-invasive recording of neuronal network function^27^.

A widely used 2D model to study neuronal function are *Neurogenin-2* (*Ngn2*) induced glutamatergic neurons^28^. Their rapid differentiation and synapse formation make them well-suited for activity assays and disease phenotyping^27,29^. Here, we present a fast and robust hiPSC-based protocol that allows for the co-maturation of microglia and with *Ngn2*-induced neurons. This manuscript describes the full optimization process and offers practical guidelines. Our co-maturation protocol enables sustained interaction throughout development and long-term co-culturing of both cell types while preserving cell type identities. The protocol promotes characteristic microglia-neuron interactions, induces the typical ramified microglia morphology and expression of microglial markers, underscoring the need for the presence of neurons to study microglial function.

## Results

We developed a co-maturation model where both hiPSC-derived microglia (iMGs) and *Ngn2*-induced neurons^28,30^ differentiated alongside each other (Fig. 1a). In our model, iMGs were added to the neurons at days *in vitro (*DIV) 9, since this is the timepoint where neuronal lineage induction was stable and doxycycline could be withdrawn to minimize side effects on microglia differentiation. From DIV 9 to DIV 28, iMGs and neurons co-matured together and developed their characteristic ramifications and dendrites, respectively (Fig. 1a). The iMGs integrated into the dense neuronal networks and formed contact sites with the dendrites. We tested the protocol using six iPSC lines for iMG differentiation across researchers and laboratories (Suppl. Fig. 1) to ensure generalizability and robustness of the protocol. In addition to a comprehensive description of the co-maturation protocol in the methods section, recommendations and troubleshooting tips can be found in Suppl. Table 1. Below, we describe in detail the steps and considerations taken to achieve this co-maturation protocol.

**Figure 1.**
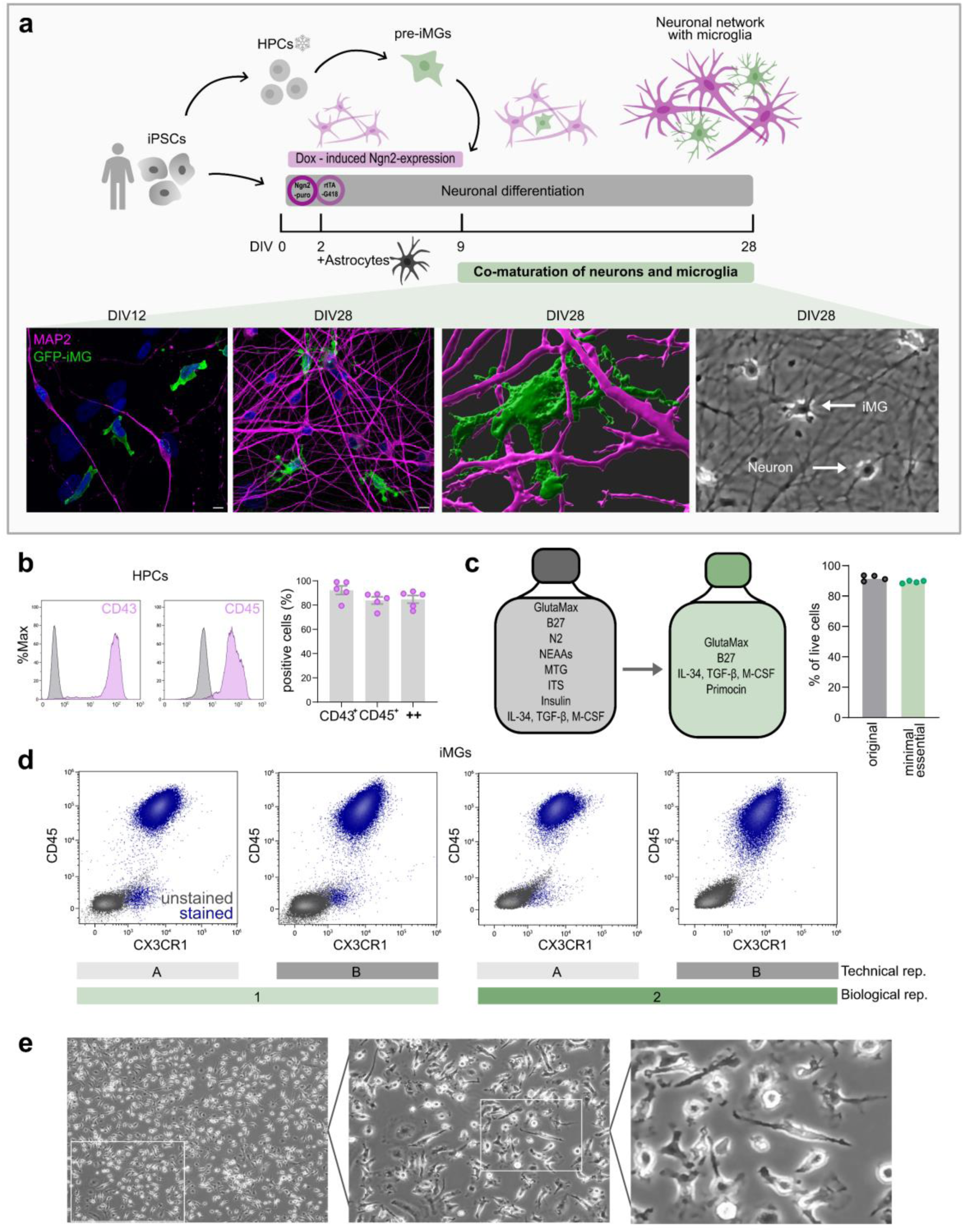
HiPSC-based protocol allowing for co-maturation of microglia and neuronal networks on the basis of a minimal essential medium for iMG differentiation. (a) Schematic protocol overview for co-maturation cultures of hiPSC-derived microglia (iMGs) and neurons (top) and representative fluorescence microscope images showing developing iMGs (green) and neurons (magenta) at DIV 12 and DIV 28 (bottom left) and DIV 28, scale bars = 10 μm, and digital reconstruction and brightfield image of iMG in neuronal network at DIV 28 (bottom right). (b) Flow cytometry of hematopoietic progenitor cells (HPCs) at DIV 11 showing representative CD43 and CD45 expression (left) and quantification of percentage of positive cells, N = 5 (right). (c) Medium composition of identified minimal essential medium for iMG differentiation (left) and live cell count comparison of medium compositions using flow cytometry, n = 4 (right). (d) Flow cytometry staining of CD45 and CXCR1 in iMGs at DIV 23, N = 2, n = 2. (e) Representative brightfield microscope image of iMGs at DIV 23.

### Defining the minimal essential medium for microglia differentiation compatible with neuronal differentiation

Co-maturation of iMGs and neurons required the identification of a medium composition that supports microglial lineage commitment while remaining compatible with *Ngn2-*induced neuronal differentiation. We began by adapting the protocol established by McQuade et al.^31^, which involves the stepwise differentiation of iMGs via hematopoietic progenitor cells (HPCs). We generated HPCs and confirmed their identity by detecting the expression of canonical markers CD43 and CD45 using flow cytometry (Fig. 1b).

To optimize HPC harvesting, we analyzed the cultures from DIV 8 to DIV 12 (Suppl. Fig. 2a). We measured the expression of CD206, a marker of differentiation into brain-associated macrophages (BAMs), which microglial precursors do not express^32^. The percentage of CD206 expressing HPCs increased from 11% at DIV 11 to 20% at DIV 12 (Suppl. Fig. 2b). To reduce variability across cells and batches, we restrained from multiple harvest days as described in the original protocol and selected DIV 11 as the standardized harvesting timepoint. Additionally, we successfully introduced a cryopreservation step to enable batch-wise HPC production, minimize variability, and streamline downstream workflows.

Next, we screened medium components using an all-minus-one fashion to identify a minimal essential medium for iMG differentiation from the HPC stage. We identified a simplified version of the original McQuade medium omitting N2, NEAAs, MTG, ITS, and Insulin (Fig. 1c, Suppl. Fig. 3a, see Methods). Besides the essential microglial growth cytokines IL-34, TGFβ, M-CSF, the remaining components are shared with the *Ngn2*-induced neuronal differentiation^30^. This minimal essential medium preserved cell viability (Fig. 1c) and maintained key marker expression (Suppl. Fig. 3). Using flow cytometry, we consistently detected CD45 and CX3CR1 co-expression in iMGs across both technical replicates (different wells from the same differentiation, denoted by n in all figure legends) and biological replicates (separate differentiations, denoted by N in all figure legends) (Fig. 1d). The iMGs exhibited typical monoculture morphology (Fig. 1e), with heterogenous shapes ranging from rod-like to slightly ramified or rounded forms.

### Minimal essential medium induces gene, protein and functional profile of microglia

We next examined the molecular phenotypes induced by our neuron-compatible iMG-differentiation. To do this, we compared the transcriptomes of iMGs differentiated in the new minimal essential medium to human post-mortem microglia (cultured or non-cultured), monocyte-derived macrophages, alveolar macrophages, iPSCs, and iPSC-derived neurons. To benchmark in comparison to existing approaches, we differentiated iMGs using the Stemcell Technologies kit, which builds on McQuade^31^. Principal component analysis showed that iMGs clustered with other microglia types and away from iPSCs, indicating that the minimal essential medium successfully induces a microglia-like transcriptional profile (Fig. 2a). However, when we focused on the myeloid cells we found that monoculture iMGs, whether differentiated with our new minimal medium or the Stemcell Technologies kit, remained distinct from postmortem microglia (Fig. 2a).

**Figure 2.**
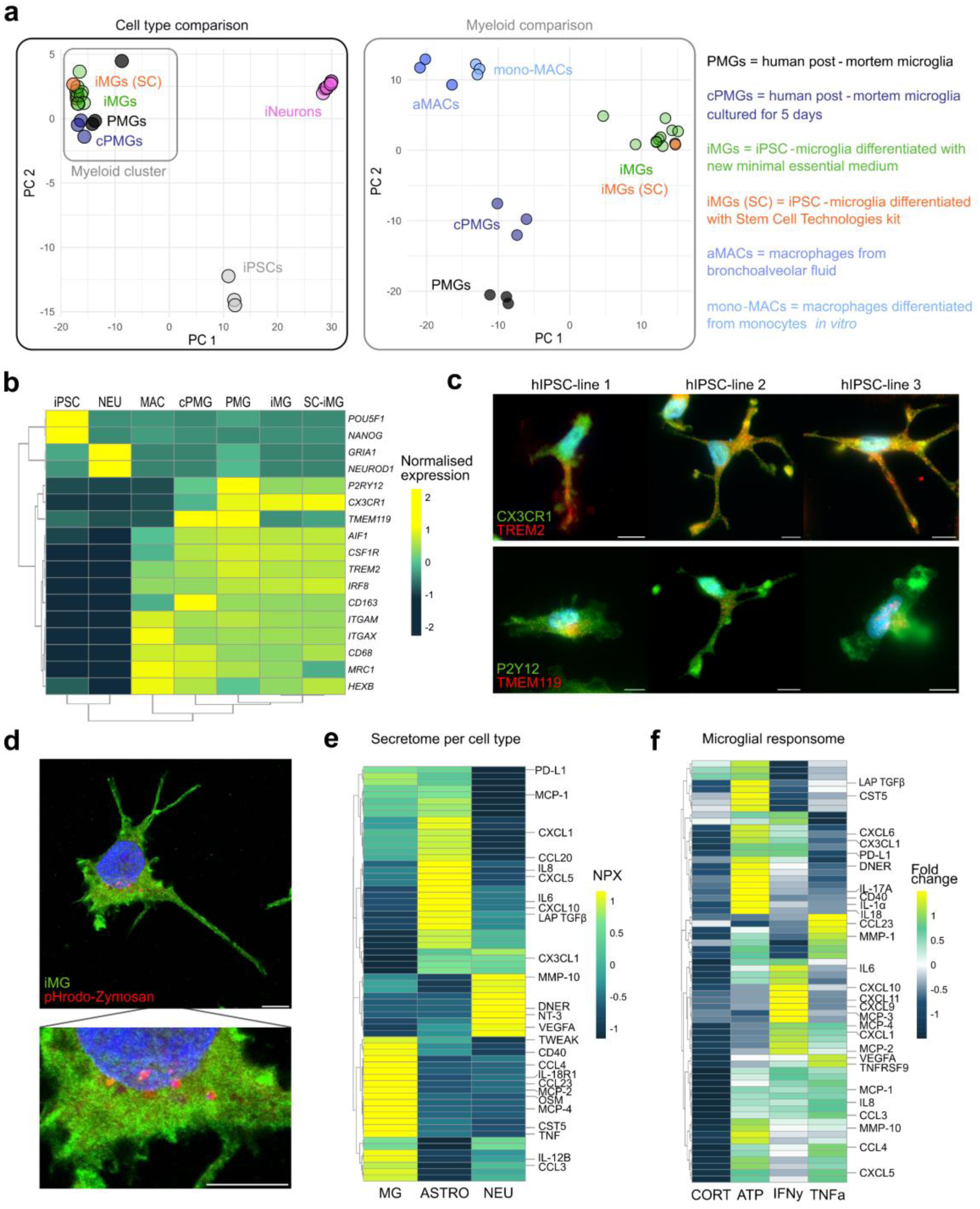
Transcriptional comparison and functional assessment of the iMGs differentiated with minimal essential medium. **(a)** Principal component analysis (PCA) of gene expression profiles of different cell types, N = 3 – 10 per cell type. **(b)** Heatmap showing average gene expression of key markers per cell type. **(c)** Representative fluorescence microscopy images of CX3CR1, TREM2, P2RY12, and TMEM119 expression in iMGs at DIV 23, scale bars = 10 μm. **(d)** Representative fluorescence microscopy image of pHrodo-labelled Zymosan bioparticles phagocytosed by iMG at DIV 23, scale bars = 10 μm. **(e)** Heatmap of secreted protein profile by iMGs (MG), astrocytes (ASTRO) and neurons (NEU), measured in monocultures, n = 3 per cell type. **(f)** Heatmap of secreted protein profile of iMGs in response to hydrocortisone (CORT), ATP, IFNγ, and TNFα, n = 2.

We then examined expression of key genes to assess the microglial identity. We found that iMGs expressed *CX3CR1*, *AIF1*, *CSF1R*, and *TREM2* at levels comparable of post-mortem microglia, but showed reduced expression of *P2RY12* and *TMEM119* (Fig. 2b). Notably, *P2RY12* expression was also reduced in cultured post-mortem microglia (cPMGs) (Fig. 2b), consistent with previous findings that are highlighting the sensitivity of homeostatic marker expression to environmental context^23^. These observations support the notion that monoculture conditions fail to induce a fully homeostatic microglial gene signature^16^ and emphasize the importance of incorporating neuronal and other CNS-resident cells, to study microglia *in vitro.* Because transcript levels do not always correspond to protein abundance^18,33^, we verified the protein levels of key proteins CX3CR1, TREM2, and P2RY12 using immunocytochemistry (Fig. 2c). As expected, based on the previous transcriptomic analysis, TMEM119 levels were low in iMGs in monoculture.

To validate microglial functionality, we assessed phagocytosis and confirmed internalization of pHrodo-labelled particles (Fig. 2d). We further tested the iMGs’ ability to respond to inflammatory stimuli by analyzing protein secretion profiles in response to stimulation with hydrocortisone, ATP, IFNγ, and TNFα using Olink proteomics. This analysis revealed cell-type specific baseline signatures and stimulus-specific protein responses (Fig. 2e-f). Together, these findings validated that our minimal essential medium induces characteristic gene expression, protein profiles and functional behaviors of microglia. We concluded that this formulation yields a robust monoculture platform comparable to established iMG differentiation protocols and offered a strong basis for the subsequent co-maturation protocol with neurons.

### Astrocytes are required for long-term microglia survival and integration into neuronal networks

After optimizing the minimal-essential medium for iMGs in monoculture, we proceeded to develop the co-maturation protocol. To support neuronal differentiation, we supplemented the medium with growth factors NT3 and BDNF. We replaced DMEM/F12 with Neurobasal, which is optimized for neuronal health and activity, while retaining all other components from the minimal essential iMG medium (see Methods). To facilitate monitoring of microglia in various conditions, we used a GFP-actin tagged hiPSC line to generate fluorescently labeled iMGs (Fig 3a-b).

**Figure 3.**
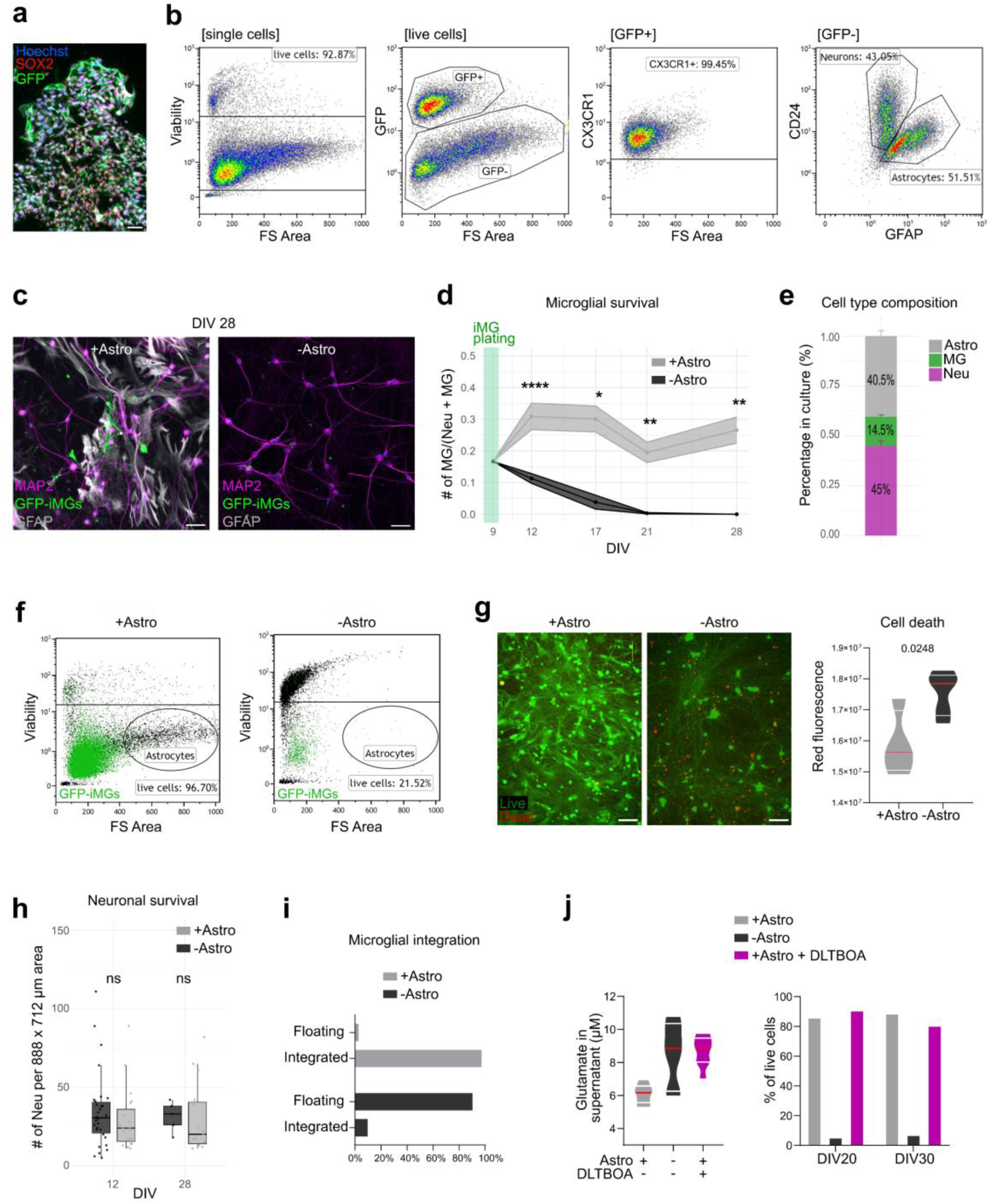
Astrocytes are required for long-term survival and integration into neuronal networks. (a) Representative fluorescence microscope image of GFP-tagged hiPSC colony, scale bar = 50 μm. (b) Flow cytometry panels showing gating strategy for the detection of GFP-positive microglia in co-cultures. **(c)** Representative fluorescence microscope images of cultures with (+Astro) or without (-Astro) astrocytes at DIV 28, scale bars = 50 μm. **(d)** Quantification of iMG numbers in cultures across development, N = 2, n = 2,102 neurons for +Astro, n = 1,854 neurons for -Astro, data represents means ±SEM, *p<0.05, **p<0.01, ****p<0.0001, unpaired t-test with post hoc Bonferroni correction. **(e)** Cell type composition in cultures at DIV 28 quantified by immunocytochemistry images, N = 2, n = 932 cells, data represents means ±SEM. **(f)** Flow cytometry panels showing GFP-iMGs and live cell percentages in co-cultures with or without astrocytes. **(g)** Representative fluorescence images of Live-Dead staining in co-cultures with or without astrocytes, scale bars = 100 μm (left) and quantification of cell death at DIV 22 using the Integrated Density of the Red Fluorescence, N = 1, n = 4-5 wells per condition, unpaired t-test, (right). **(h)** Number of neurons per 20x microscopy image in co-cultures with or without astrocytes quantified at DIV 12 and DIV 28, N = 2, unpaired t-test. **(i)** Percentage of microglia integrated in culture vs floating in supernatant at DIV 20 measured by flow cytometry, N = 1. **(j)** Concentration of Glutamate (μM) in supernatant of cultures across DIV 14 – DIV 20, N = 1, n = 8 (left) and the percentage of live cells in tricultures measured by flow cytometry at DIV 20 and 30, N = 1 (right).

During protocol optimization, we identified astrocytes as essential for establishing a stable long-term co-culture. Within the neuron field, it is standard practice to add rodent *ex vivo* astrocytes to promote healthy neuronal differentiation and function *in vitro*^28,30,34^. When we omitted astrocytes at DIV 2, microglial numbers declined rapidly and were undetectable by DIV 21 (Fig. 3c-d). In contrast, when we added astrocytes, microglial numbers remained stable from DIV 10 to DIV 28 (Fig. 3d). While most of our experiments were performed at DIV 28, we could maintain the co-maturation culture until DIV 51 (Suppl. Fig. 1). We characterized the cell-type composition at DIV 28 using immunocytochemistry. Because techniques such as flow cytometry and single-cell RNA sequencing introduces biases, due to the difficulty in dissociating dense neuronal networks, we opted for imaging-based quantification (Fig. 3c). Neurons comprised approximately 45% of the cultures, microglia 15%, and astrocytes the remaining 40% (Fig. 3e).

We hypothesized that the microglia loss in the absence of astrocytes resulted from either impaired integration into the network or increased cell death. To test this, we performed flow cytometry and observed a drastic reduction of live cell counts in the condition without astrocytes (96% to 21%) accompanied by a reduction of GFP-iMGs (Fig. 3f). Additional live cell imaging confirmed increased cell death when astrocytes were not present (Fig. 3g). Since neuronal numbers remained stable from DIV 12 to DIV 28 when cultures without astrocytes (Fig. 3h), the observed cell death could be attributed to the microglia.

Next, we quantified microglial integration by comparing microglial numbers in the culture supernatant and the adherent fraction after dissociation using flow cytometry. The presence of astrocytes increased microglial integration from ∼10% to 97% (Fig. 3i), indicating that astrocytes promote both microglial survival and integration. We speculated that toxic glutamate accumulation in neuronal networks due to the lack of astrocytic clearance, could be driving this effect. Indeed, glutamate levels were higher in the supernatant of co-cultures without astrocytes (Fig. 3j). We pharmacologically blocked excitatory amino acid transporters (EAATs) with DL-TBOA in co-cultures with astrocytes leading to an increase of glutamate to the level of the - Astro condition (Fig. 3j). However, treatment with DL-TBOA did not mimic the decrease in viability observed in co-cultures without astrocytes (Fig. 3j), suggesting that alternative mechanisms underlie the astrocyte-dependent support.

### Co-maturation cultures show cell-type specific profiles and mature neurons with organized synaptic activity

To fully characterize the co-maturation cultures on the transcriptome level, we performed single-cell RNA sequencing on networks with hiPSC-derived iMGs and neurons complemented with rodent astrocytes at DIV 28. Clustering with the Louvain algorithm after Harmony integration revealed three distinct clusters (Fig. 4a). Based on signature marker expression of the respective cell types, we annotated microglia, neurons, and astrocytes (Fig. 4b). All batches, prepared in separate libraries, showed the presence of all three cell types, and cultures without addition of iMGs contained no cells clustering to the microglia cluster (Fig. 4a). The neuronal cluster expressed synaptic markers like *SYP, SYT1, and SLC17A6* indicating lineage commitment to excitatory glutamatergic neurons (Fig. 4c). We detected no pluripotent marker expression in any of the three clusters, confirming successful differentiation with no remaining hiPSC colonies in the cultures (Fig. 4c). Next, we verified the expression of Synapsin1 using immunocytochemistry, indicating the presence of synapses on the neurons (Fig. 4d). To validate the functional neuronal network phenotype of the co-maturation cultures, we recorded spontaneous activity using multi-electrode arrays (MEA) over time (Fig. 4e). We observed an increase in general spiking activity, reflected by the mean firing rate, from DIV 14 to 28 and the synchronization of network activity, reflected by establishment and increase in network burst percentage after DIV14 (Fig. 4f). Additionally, on single neuron level we verified functional synaptic connectivity in the co-maturation cultures using whole-cell patch clamp recordings, both by measuring miniature excitatory postsynaptic currents (Fig. 4g) and by optogenetically inducing synaptic responses in presynaptic channelrhodopsin2 (ChRh2) expressing neurons (Fig. 4h, Suppl. Fig. 4). Taken together, the co-maturation cultures show cell-type specific transcriptomic profiles and functional properties of neuronal activity on single-cell and network level.

**Figure 4.**
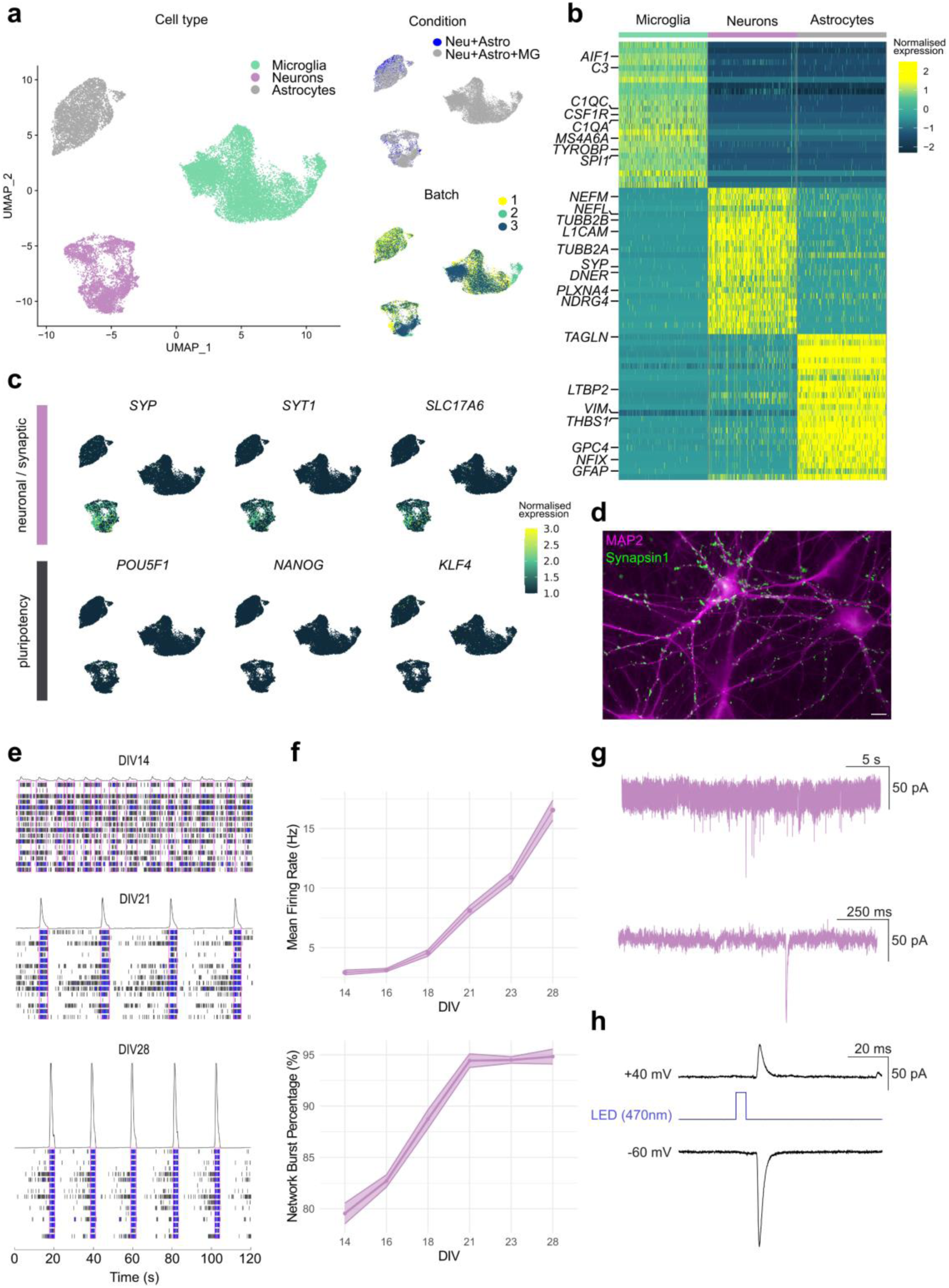
Transcriptional and functional validation of co-maturation cultures. **(a)** Uniform manifold approximation and projection (UMAP) analysis of single-cell RNA-seq data of cultures at DIV 28 showing clustering of the three cell types (left) and distribution of condition and batch (right), N = 3, n = 24,791 single cells. **(b)** Heatmap showing expression profile of the top 25 genes per cell-type. **(c)** UMAP analysis showing expression of neuronal (top) and pluripotent (bottom) markers. **(d)** Representative fluorescence microscopy image of Synapsin1 (green) on neuronal dendrites (MAP2, magenta), scale bar = 10. **(e)** Representative multi-electrode array (MEA) raster plots with network bursts annotated in magenta showing increased organization of neuronal network activity from DIV14 to DIV 28. **(f)** Quantification of mean firing rate (Hz; top) and network burst percentage (%; bottom) across 6 developmental timepoints, N = 4, n = 81 neuronal networks. **(g)** Representative traces of miniature excitatory postsynaptic currents (mEPSCs) recorded from single neurons at DIV 49 using whole-cell patch clamp recordings. **(h)** Traces of light-evoked synaptic currents recorded from a channelrhodopsin-2-negative neuron at DIV 28.

### Co-maturation protocol induces signature morphology, transcriptome and proteins of human microglia

Microglia undergo rapid changes in gene expression, protein levels and morphology when they are removed from the CNS^18,22^. Vice versa, co-culture and xenotransplant models have shown that the interaction with CNS-resident cells induces typical microglia features^23,24^. Given the continuous interactions between iMGs and neurons in our culture model, we speculated that the co-maturation approach may support microglia identity. We therefore assessed the impact of our co-culture on iMG morphology, gene and protein expression. First, we examined iMG ramification in the neuron-glia networks (co-culture) compared to iMG monocultures. Immunocytochemistry analysis of GFP-tagged iMGs showed a significant increase in ramification in the co-culture condition (Fig. 5a). We observed high levels of morphological heterogeneity in both conditions, but a robust effect on ramification by the presence of neuron-glia networks (see representative microscopy images in Fig. 5a).

**Figure 5.**
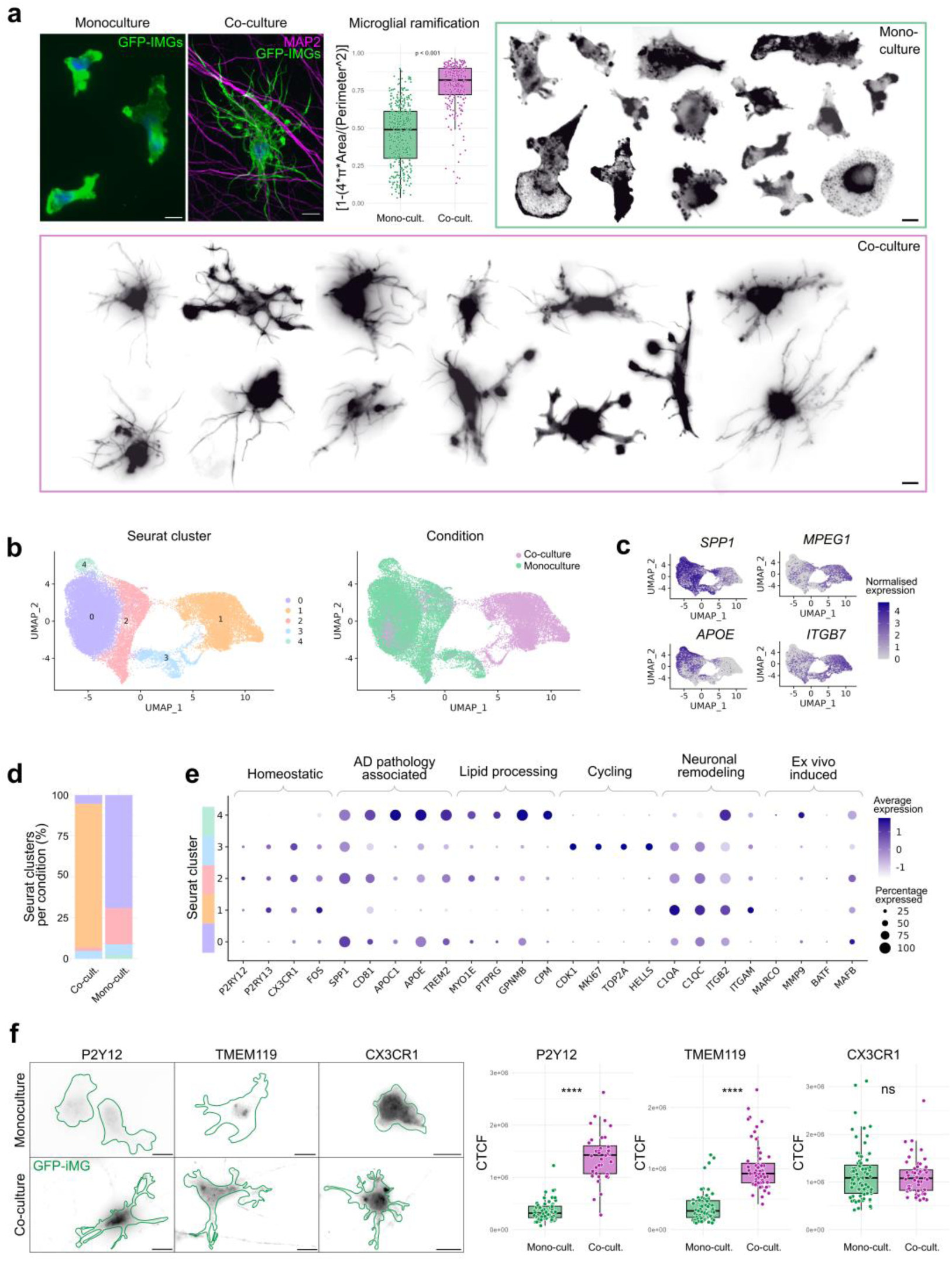
Morphological, transcriptomic, and proteomic changes of iMGs in the co-maturation protocol compared to monoculture. **(a)** Representative fluorescence microscope images of GFP-actin based iMG morphology in monoculture vs. the co-culture with neurons at DIV 28, scale bars = 10 μm (top left), quantification of ramification index per cell, N = 3, n = 352 for monoculture, n = 249 for co-culture, unpaired t-test (top middle), and inverse LUT images of iMGs per culture condition, scale bars = 10 μm (top right and bottom). **(b)** UMAP analysis of single-cell RNA seq data of iMGs at DIV 28 showing Seurat clusters (left) and the culture condition (right), N = 3, n = 42,226 single cells. **(c)** UMAP analysis showing top differentially expressed genes of co-culture vs. monoculture iMGs (Average Log2 fold changes are: *SPP1* = -4.35, *APOE* = -4.29, *MPEG1* = 2.99, *ITGB7* = 2.9, all show p<0.0001). **(d)** Distribution of Seurat clusters per culture condition. **(e)** Average gene expression per Seurat cluster highlighting signatures associated with different microglia contexts. **(f)** Representative fluorescence microscope images of P2RY12, TMEM119, and CX3CR1 protein expression in iMGs in monoculture vs co-culture, cell outline determined by GFP-actin signal, scale bars = 10 μm (left) and quantification of fluorescence signal per cell, CTCF = corrected total cell fluorescence, N = 2, P2RY12; n = 59 for monoculture, n = 36 for co-culture, TMEM119; n = 68 for monoculture, n = 57 for co-culture, CX3CR1; n = 80 for monoculture, n = 55 for co-culture, unpaired t-test (right).

To assess the effect of the co-maturation at the transcriptomic level, we compared iMGs from both culture conditions using single-cell RNA sequencing. In this experiment, the initial starting vials of HPCs, coating, medium master mixes, and cell culture schedule were identical to ensure the presence of neuron-glia networks is the only variable factor. Unsupervised clustering revealed a clear distinction between the two conditions (Fig. 5b) indicating that the neuron-glia network environment substantially changed gene expression in iMGs. Using differential analysis, we identified *SPP1* and *APOE* as the top downregulated genes in co-culture iMGs compared to monoculture iMGs (log fold change below -4). Interestingly, these are the top two contributors to the disease-associated microglia signature often observed in Alzheimer’s disease (AD) pathology^35^. Among the top upregulated genes in co-culture iMGs were *MPEG1* and *ITGB7*, which is coding for an integrin protein involved in cell-cell adhesion (Fig. 5c). Next, we investigated characteristic gene sets of microglia substates and functions based on transcriptional studies of human postmortem microglia^22,36–38^, the environment-dependent signature described by Gosselin et al.^16^, and the current consensus in microglia nomenclature^8^. Cluster 1, where the majority of co-culture iMGs were distributed in (Fig. 5d), showed an upregulation of homeostatic and neuronal remodeling markers (Fig. 5e). In contrast, genes associated with AD pathology and lipid processing were strongly downregulated compared to all other clusters. The most prominent cluster in the monoculture iMGs (cluster 0) showed low expression for the homeostatic genes. Cluster 3 was identified as cycling microglia and contained both monoculture and co-culture iMGs. Cluster 4 comprises a small subset of monoculture iMGs with highest expression of AD pathology related genes and no homeostatic signature gene expression (Fig. 5e). Genes associated with the *ex vivo* signature induced by culturing of post-mortem microglia^16^ were expressed lower in co-culture iMGs (cluster 1) compared to monoculture iMGs (cluster 0, 2, 4) (Fig. 5e). This suggests that the co-culture environment partially buffers the effects observed when microglia are cultured without brain context.

Next, we assessed a panel of key microglia markers on protein level using immunocytochemistry (Fig. 5f). Analysis of the fluorescence per cell revealed significant upregulation of P2RY12 and TMEM119 in co-culture iMGs (Fig. 5f). CX3CR1 expression remained unchanged, consistent with its already elevated mRNA levels in monoculture iMGs (Fig. 2b). In conclusion, the neuron-glia network in our co-maturation model promotes several aspects of *in vivo* microglia identity, highlighting the importance of using long-term co-cultures protocols to model microglia *in vitro*.

### Co-maturation cultures show key features of microglia-neuron crosstalk

*In vivo* research suggests interaction of microglia with key sites like synapses^6,7^ or the axon initial segment (AIS)^39^. We confirmed these characteristic neuron-microglia interactions in our *in vitro* co*-*maturation model using immunocytochemistry. iMGs formed contact sites with dendrites and axons, as well as postsynaptic marker Homer1 and the AIS (Fig. 6a-b). Live cell imaging further showed continuous and rapid movement of microglial processes within the neuron-glia network and interaction with neuronal branches (Fig. 6c, Suppl. Video 1).

**Figure 6.**
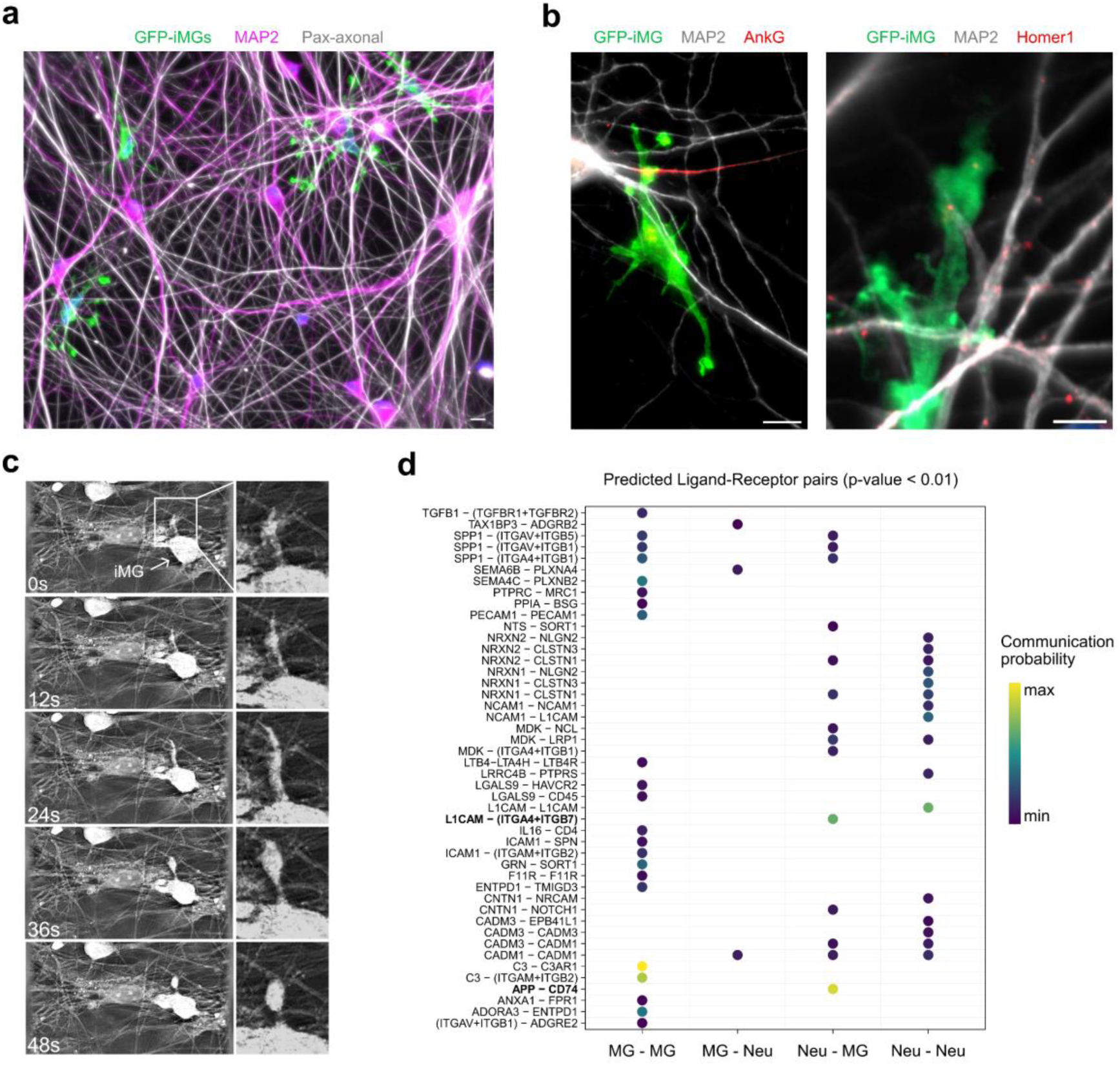
Co-maturation protocol allows for dynamic microglia-neuron interactions at dendrite-, axon-, and synapse level. **(a)** Representative fluorescence microscope images of microglial interaction with neuronal dendrites (MAP2) and axons (pan-axonal) at DIV 28, scale bar = 10 μm. **(b)** Representative fluorescence microscope images of microglial interaction with axon initial segment (AnkG), scalebar = 10 μm (left) and post-synapse (Homer1), scale bar = 5 μm (right) at DIV 28. **(c)** Refractive index live-cell imaging timeline showing microglial process interacting with neuron at DIV 28. **(d)** Predicted ligand-receptor pairs of microglia and neurons in co-maturation cultures based on single-cell RNA sequencing data at DIV 28, N = 3, n = 24,791 single cells.

Next, we explored receptor-ligand interactions using the CellChat algorithm^40^ in our single-cell transcriptomic dataset (Fig. 6d). We observed expected microglial communication pathways like TGFβ- or complement (C3) signaling. For crosstalk between neurons and iMGs the top predicted interactions were *L1CAM – (ITGA4 + ITGB7) and APP – CD74.* Interestingly, *ITGB7* was among the top upregulated genes in co-culture iMGs compared to monoculture iMGs (Fig. 5c) and this upregulation might be driven by interaction with neuronal adhesion molecules. These findings indicate that the long-term duration of our co-maturation approach allows for dynamic and extensive neuron-microglia interactions.

### Human iPSC-derived astrocytes are compatible with the long-term iMG-neuron co-maturation protocol

To engineer a fully hiPSC-derived triculture, we successfully implemented our co-maturation protocol with human iPSC-derived astrocytes (iAs). The astrocyte part of the protocol is based on our previously optimized protocol^41^. As we developed those protocols alongside each other to achieve compatibility, they follow similar strategies. The only modification to the rodent astrocyte co-maturation protocol was the introduction of human astrocytes at DIV 6 instead of DIV 2 (Fig. 7a). As described before, iMGs were added after doxycycline withdrawal at DIV 9.

**Figure 7.**
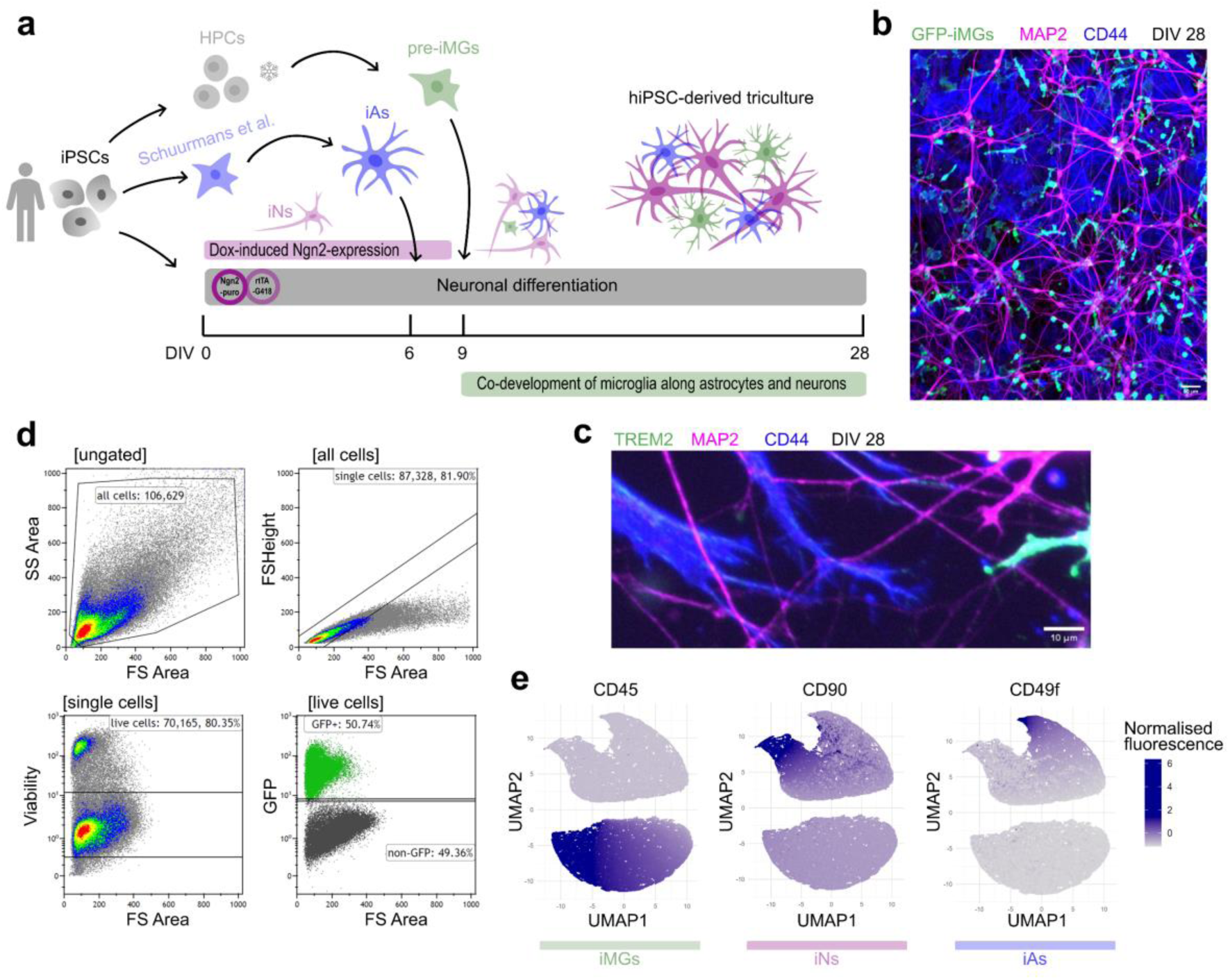
Human iPSC-derived astrocytes are compatible with iMG-neuron co-maturation protocol and also support microglial survival. **(a)** Schematic protocol overview for generation of tricultures of hiPSC-derived astrocytes (iAs), microglia and neurons. **(b)** Representative fluorescence microscope images showing iAs (CD44), iMGs (GFP) and neurons (MAP2) at DIV 28, scale bar = 20 μm. **(c)** Representative fluorescence microscope images showing processes of iAs (CD44), iMGs (GFP) and neurons (MAP2) at DIV 28, scale bar = 10 μm. **(d)** Flow cytometry of hiPSC-derived tricultures showing gating strategy and presence of GFP-positive cells at DIV 28. **(e)** Uniform manifold approximation and projection (UMAP) analysis of CD45, CD90 and CD49f flow cytometry signal.

At DIV 28, we could still detect the GFP-tagged iMGs using both immunocytochemistry (Fig. 7b-c) and flow cytometry (Fig. 7d), confirming that the iAs were able to promote integration and long-term survival of iMGs. Furthermore, this showed that microglial survival was not driven by the *ex vivo* nature of the rodent astrocytes. Both iAs and iMGs contacted neuronal dendrites with their protrusions (Fig. 7c). Using flow cytometry on the fully hiPSC-based tricultures we reliably separated iMGs, neurons, and iAs with unsupervised clustering of surface marker expression of CD45, CD90, and CD49f, confirming successful differentiation into all three respective cell types (Fig. 7e).

## Discussion

Microglia are highly sensitive to their surroundings. This environment-dependent transcriptional and functional signature makes their *in vitro* modeling inherently challenging^16,22^. Consequently, monoculture systems of both primary and hiPSC-derived microglia are lacking essential CNS-derived signals required to recapitulate their physiological characteristics^20,21,23^. Neurons and microglia continuously receive signals from each other influencing maturation and function. Thus, models used to understand neuron-microglia communication should reflect this ongoing interplay between microglia, neurons and other CNS-resident cells like astrocytes. To address this, we developed a hiPSC-based long-term culture model, in which microglia and neurons co-mature. Our protocol offers a robust and efficient platform to comprehensively study human neuro-immune crosstalk.

The continuous interaction during co-maturation induced characteristic microglia-neuron interactions and improved the microglia signature on morphology, gene, and protein level. The presence of the hiPSC-derived neuron-glia networks led to pronounced microglial ramification similar to how microglia are observed in xenotransplanted organoids^23^. This characteristic microglial morphology is accompanied by an upregulation of markers associated to neuronal remodeling and homeostatic microglia^8,22^, highlighting the importance of contact to other CNS-resident cells for establishing microglia identity *in vitro*. Interestingly, genes and signaling pathways associated with neurodegenerative pathology were strongly downregulated in microglia in the co-cultures compared to monocultures. These included *APOE and SPP1,* which have been widely studied for their role in AD^16,35,38,42,43^. Our findings may suggest that a loss of healthy signaling from neurons might drive the characteristic disease-associated phenotype both in microglia from AD patients and in microglia monocultures. This further highlights that our co-maturation model could mitigate the stress typically induced by culturing microglia *in vitro*, which normally skews them toward a neurodegenerative-associated substate. As a result, our co-culture provides a suitable platform for studying microglial roles in a variety of contexts.

During optimization of our co-maturation model, we discovered a fundamental role of astrocytes for microglial survival and integration. Several studies have highlighted microglia-astrocyte crosstalk in the past^44^. Especially the astrocytic secretion of growth factors has been shown to be critical for microglia differentiation^45^. However, it is likely that the supportive function of astrocytes extends beyond the supply of trophic factors like TGF-β and M-CSF^46^, as we supplemented both monocultures and co-cultures with these factors (see Methods). Moreover, the protective effect of astrocytes might also be mediated indirectly via effects on the neurons. The survival of microglia in monoculture, but not in neuron-only co-cultures, suggests that neurons may release stress factors when astrocytes are absent. Astrocytes promote neuronal growth^34^ and synaptic maturation^28^ in cultured neurons, and are therefore widely used for *in vitro* studies of neuronal function^30^. While neuronal numbers remained stable when cultured without astrocytes, their health and maturity may have been compromised. This could in turn limit microglial health and integration into the neuronal networks. Next to rodent astrocytes, we also streamlined this co-maturation protocol for use with our hiPSC-derived astrocyte protocol^41^. This allows for the fast and robust generation of a fully hiPSC-derived triculture with neurons, astrocytes, and microglia, expanding the possibilities in studying human brain function.

Advances in iPSC technologies have yielded complex 3D models like organoids for *in vitro* studies. While these models have advantages depending on the research question, especially for understanding cortical development, they are challenging to scale up for high-throughput screening. The 2D format enhances scalability and facilitates applications like (live-cell) imaging, micro-electrode arrays, and drug screening. This protocol allows for using cost-effective high-throughput pipelines while maintaining sufficient complexity to model cell-cell interactions and functional phenotypes that reflect physiological conditions. Furthermore, our co-maturation approach allows for full control of the genotype of microglia, neurons, and astrocytes posing it especially useful to disentangle cell-type specific contributions to disorders with a known genetic component. We provide troubleshooting tips for our co-maturation approach (e.g. working with disease lines, detachment issues, remaining hiPSC colonies, clustering) in Suppl. Table 1. Variations of the co-maturation protocol to accommodate specific requirements can be made (e.g. human or rodent astrocytes, addition of inhibitory neurons^47^). Our improved co-maturation model of hiPSC-derived microglia with neuron-glia networks provides a powerful tool to investigate neuro-immune crosstalk in health and disease, and specifically elucidate the role of microglia in modulating neuronal function and development.

## Methods

### Human induced pluripotent stem cell lines and maintenance

Quality controlled human induced pluripotent stem cell lines (hiPSCs) were used. These include WTC-11 (Coriell Institute, GM25256) with an endogenous actin-GFP tag (Allen Institute, AICS-0016 cl.184), HPSI0314i-hoik_1, and HPSI0114i-kolf_2 (Human Induced Pluripotent Stem Cells Initiative, Wellcome Trust Sanger Institute). All three of these hiPSC lines are fibroblast derived. WTC-11 underwent episomal reprogramming, and the HPSI lines were reprogrammed with non-integrating Sendai virus.

hiPSCs were cultured in Essential 8^TM^ Flex Basal Medium (Gibco, A2858501) supplemented with Primocin (0.1 μg/ml, Invivogen; ANT-PM-2) on Geltrex-coated (Gibco, A1413301) 6-well plates (Corning, 353046) at 37 °C/5% CO2. Medium was refreshed every 2-3 days. At around 80% confluency the hiPSCs were passaged using the enzyme-free reagent ReLeSR (Stemcell Technologies, 100-0483). Passaging was done with wide-tip pipettes to keep hiPSC colonies intact and reduce stress. All hiPSCs were tested biweekly for mycoplasma contamination using MycoAlert PLUS (Lonza, LT07-703).

### iMicroglia differentiation

The differentiation of hiPSC-derived microglia (iMGs) follows a two-step protocol with initial hematopoietic progenitor cell (HPC) generation (Suppl. Fig. 2a), an optional freezing step for streamlining, and subsequent microglia differentiation. HPCs were generated with the HPC Kit from Stemcell Technologies according to the guidelines provided if not stated otherwise (05310). Assessing marker expression over the timeline of differentiation revealed that HPCs are best harvested at DIV 11 (Suppl. Fig. 2). We restrained for multiple DIV harvest to ensure overall quality and reduce variability in differentiations. During our optimization we found that HPC quality determines the success of the microglia differentiation, and we therefore only used harvests that showed >95% of CD43 and CD45 positive cells. The HPCs were frozen in a Mr. Frosty Freezing Container (Thermo Fisher) using PSC Cryomedium (Gibco, A2644601) and stored in liquid nitrogen at 500k/vial until further use.

HPCs were thawed in minimal essential microglia medium. This medium consists of DMEM/F12 base (Gibco, 11320074) supplemented with GlutaMAX (1x, ThermoFisher, 35050061), B27 (1x, Gibco, 17504044), Primocin (100 μg/ml, Invivogen, ant-pm-2), IL34 (100 ng/ml, Peprotech, 200-34), TGF-b (50 ng/ml, Peprotech, 167100-21), M-CSF (25 ng/ml, Peprotech, 167300-25). To maximize cell survival, thawing must be conducted fast. Standard pipette tips can be used due to the single-cell nature of HPCs. HPCs were seeded onto six-well plates coated with Geltrex (1:100, Gibco, A1413202) at a density of 250,000 HPCs per well. We found that a lower starting density is not beneficial for microglia differentiation. Throughout the differentiation process, iMG cultures were refreshed 2-3 times a week by adding one milliliter per well in a six-well plate without media aspiration to retain floating microglia. After around 5 days, the wells became highly confluent and the iMGs were split by transferring the floating iMGs to new TC-treated wells that do not require additional coating for microglial attachment after the first 3-5 days. In this first split, we supported proliferation and survival by addition of GM-CSF (10 ng/ml, Peprotech, 300-03). However, during the following splits and the subsequent culture process, we restrained using it to not compromise on microglia phenotype.

During culturing the next weeks, microglial cultures were split upon the observation of cell aggregates or medium overflowing. This typically occurred around DIV 8 and DIV 15 but depended on the cell line and batch used. Splitting strategy was optimized to maximize cell health under different circumstances. If no extreme cell aggregates were observed (e.g. the first splitting was primarily performed to prevent medium overflowing) only the floating cells were transferred to new wells, avoiding the detachment of adherent iMGs. This approach minimizes centrifugation stress, preserves ramified microglial morphology and improved differentiation outcomes. In case GM-CSF was used (first split) it was also added in equal concentration to the original well. In case the iMGs formed clumps that continued to grow, cells were dissociated with TripLE (Gibco, 12604021) using a P1000 pipette. The collected cells were centrifuged at 400 × g for 5 min, resuspended in fresh base medium with cytokines, and replated in new TC-treated wells at appropriate densities. With both splitting methods, the transferred cells attached and showed no differences in morphology or marker expression compared to the original adherent cells from the starting wells.

We observed that microglial proliferation was high in the initial ten days but declined over time. Therefore, we restrained for splitting late during the differentiation process. Low density of iMGs (>250k per well in 6-well plate) limited cell survival. Example pictures of iMG cultures and recommendations on when to split can be found in Suppl. Fig. 5. Throughout the differentiation period, cultures were monitored for morphology, and appropriate density adjustments were made to optimize iMG yield and phenotype.

### *Ngn2*-induced neuronal differentiation

We differentiated hiPSCs to excitatory cortical neurons as previously described^30^, but introduced minor adjustments to allow for microglia co-culture. HiPSCs were kept at 37°C/5%/CO2. To allow differentiation, hiPSCs were transduced with lentiviral *Ngn2* and rtTA constructs and selected with increasing concentrations of G418 (100 μg/mL - 250 μg/ml, Sigma-Aldrich, G8168) and puromycin (1 μg/mL, 2 μg/ml, Sigma; P9620) respectively. Upon successful lentiviral integration, neuronal differentiation by *Ngn2* overexpression can be initiated after seeding single cells by doxycycline treatment.

Coverslips for immunocytochemistry were treated with nitric acid for three days, washed 20 times with ultra-pure water and sterilized at 180 °C for 5 h to ensure a clean surface for attachments of the neuron. To support attachment of the neurons, the wells or coverslips were first coated with poly-L-ornithine (50 ug/ml, PLO, Sigma, 3655) diluted in borate buffer for at least 3 h. After washing with ultra-pure water without drying out the wells, the wells were incubated with human laminin (5 μg/ml, Biolamina, LN521-05) overnight at 4 °C. Induction of neuronal differentiation (DIV 0) was done by dissociating hiPSC colonies into single cells using TrypLE and seeding onto the pre-coated wells in E8 basal medium (Gibco; A1517001) supplemented with RevitaCell™ (1:100, Gibco, A2644501), doxycycline (4 μg/ml, Sigma, D5207), and Primocin (100 μg/ml, Invivogen, ant-pm-2). On MEA-chips, droplet-plating was performed to ensure successful coverage of the electrodes. We plated 35k cells in 24-well plates with coverslips for immunocytochemistry, 300k cells in 6-well plates for flow cytometry and 26k cells on 48-well MEA plates (Axion Biosystem) for neuronal activity recordings. The day after plating (DIV 1) a full medium change to DMEM/F12 (Gibco, 11320074) supplemented with 1x N2 (Gibco, 17502001), 10 ng/mL NT3 (Peprotech; 450-03), 10 ng/mL BDNF (Peprotech; 450-02), 100 μM MEM non-essential amino acid solution (NEAA; Sigma-Aldrich; M7145), 4 µg/µl doxycycline, and 100 μg/ml Primocin was performed.

Neuronal maturation was supported by E18 rodent astrocytes, which were added to the culture in a 1:2 ratio at DIV 2. For the addition of hiPSC-derived astrocytes to *Ngn2*-induced neurons, please see our optimized protocol^41^. The neuronal culture component is identical to this one as both protocols are streamlined to allow for hiPSC-derived tricultures of neurons, astrocytes, and microglia. The manuscript is written in a similar style to this and provides a step-by-step guide to astrocyte differentiation including practical recommendations.

From DIV 3 onwards, the medium was switched to neurobasal medium (Gibco, 21103049) supplemented with B-27, GlutaMAX, Primocin, NT3, BDNF, and doxycycline. To remove proliferating hiPSC colonies and astrocytes from the cultures Cytosine β-D arabinofuranoside (Ara-C; 1 µM; Sigma-Aldrich; C1768) treatment was performed once at DIV 3. From DIV 6 onwards, a half medium change was performed three times a week (Mon, Wed, Fri). To prevent stress on the neurons and circumvent artifacts for later microglia addition, doxycycline was withdrawn from the medium starting at DIV 8 in all conditions. FBS was added to support astrocytic survival from DIV 9 onwards (2.5 % Sigma, F7524). Wells with unequal cell densities or uneven distribution (clustering, highway-formation, see troubleshooting tips) were excluded from analysis. The cultures were kept at 37 °C/5% CO2.

### Microglia-neuron co-maturation culture

Microglial and neuronal differentiation was initiated at the according to the guidelines described above. The microglial culture was always started on day prior to the neuronal differentiation. At DIV 9 of the neuronal differentiation corresponding to DIV 10 of microglial differentiation, they were combined. First, a half-medium change was performed to introduce the three microglial-supporting cytokines IL34, TGFβ, and M-CSF to the neuronal cultures. To avoid potential effects on the differentiating microglia, it is important that doxycycline was subsequently omitted from the medium. The complete composition of the co-maturation medium is Neurobasal medium (Gibco, 21103049) supplemented with GlutaMax (1x, ThermoFisher, 35050061), B27 (1x, Gibco, 17504044), Primocin (100 μg/ml, Invivogen, ant-pm-2), NT3 (10 ng/ml, Preprotech, 450-03), BDNF (10 ng/ml, Peprotech, 450-02), FBS (2.5%, Sigma, F7524), IL34 (100 ng/ml, Peprotech, 200-34), TGFβ (50 ng/ml, Peprotech, 167100-21), M-CSF (25 ng/ml, Peprotech, 167300-25). We found that withdrawal of FBS and consequently the formation of reactive astrocytes (based on their morphology) is more detrimental for microglial activation and survival than the addition of FBS. Therefore, we comprised with adding FBS at low dosage of 2.5%.

To not disrupt the cultures and remove secreted growth factors, it is important that the medium transition was conducted as a half-medium change. We therefore removed half the medium from the wells with the neurons, prepared the co-maturation medium with double the concentration of the cytokines IL34, TGFβ, and M-CSF and added it in a 1:1 ratio to the existing medium. This is possible because the compositions of the neuronal and co-maturation medium are identical, except for the inclusion of three cytokines. Then, the plates were incubated at 37 °C/5% CO_2_ for at least 30 min to calibrate temperature and CO_2_ levels before adding iMGs. The iMGs were harvested with TrypLE (Gibco, 120604021), centrifuged (400g) at room temperature for 5 min and resuspended in co-maturation medium to obtain a single-cell suspension. After counting with Trypan blue, iMGs were added to the neurons in a 1:5 ratio. In 24-well we plated 35k neurons, 20k astrocytes, and 7k iMGs. In 6-well plates, we plated 150k neurons, 100k astrocytes, and 30k microglia. In 48-well MEA plates (Axion Biosystems) we plated 26k neurons, 13k astrocytes, and 5.2k microglia.

The iMGs can be added into the full medium (no droplet plating needed) as they move towards the neuron-glia networks and integrate by themselves. Half of the medium was refreshed with co-maturation medium three times a week (Mon, Wed, Fri). Medium was always carefully taken from the upper edge of the well to not disrupt microglia integration into the cultures. The co-cultures were kept at 37 °C/5% CO_2_ throughout the entire differentiation process.

### Pharmacology

All reagents were prepared into concentrated stocks and stored at −20°C. For stimulations of microglia cultures, we used hydrocortisone (2.5 µM, Sigma-Aldrich, H0888), ATP (1 mM, Sigma-Aldrich, 74804-12-9), TNFα (100 ng/ml, Peprotech, 300-01A) and IFNy (100 ng/ml, Peprotech, 300-02). All concentrations were tested for potential toxicity effects and cell death before. If possible, compounds were diluted in ultra-pure water (TNFα, IFNy, ATP). Otherwise, dimethyl sulfoxide (DMSO) was used (hydrocortisone). For all pharmacological treatments and experiments, the amount of DMSO in the cell culture medium was ≤0.5% v/v.

### Immunocytochemistry

Unless stated otherwise, subsequent steps were performed at room temperature and solutions were made in 1XPBS. Cultured cells were fixed by using a gradient of 2% and subsequently 4% paraformaldehyde (PFA) supplemented with 4% sucrose. First, PFA was added to the culture in a 1:1 ratio with existing medium. After 7 min, all the medium was carefully removed from the well and 4% PFA was added for additional 7 min. We found that this process preserved cellular morphologies compared to a direct change to PFA from medium. After PFA incubation, cells were washed three times with 1XPBS, and permeabilized with 0.2% Triton (Sigma-Aldrich, T8787) for 10 min. To block nonspecific binding sites, cells were incubated with 3% bovine serum albumin (BSA, Sigma, A7906) for 1 h. Primary antibodies were diluted in the same buffer and incubated overnight at 4°C. The following primary antibodies and dilutions were used: rabbit anti-CX3CR1 (1:300, Invitrogen, PA5-19910), rat anti-TREM2 (1:300, R&D Systems, MAB17291-SP), rabbit anti-P2RY12 (1:300, Alomone Labs, APR-012), mouse anti-TMEM119 (1:300, Biolegend, A16075D), guinea pig anti-MAP2 (1:1000, Synaptic Systems, 188004), mouse anti-panAxonal (1:1000, BioLegend, 837904), rabbit anti-GFAP (1:1000; Sigma-Aldrich, AB5804), rabbit anti-Synapsin I (1:500, Sigma-Aldrich, AB1543P), mouse anti-Homer1b/c (1:200, Synaptic Systems, 160111), mouse anti-ankyrin-G (1:1000, Invitrogen, 33-800).

Secondary antibodies conjugated to either Alexa Fluor 488, 568, or 647 (Invitrogen) were incubated for 1h at 1:1000 in 1% BSA. Cell nuclei were stained with Hoechst (0.01%, ThermoFisher, H3570) for 10 min. After three final washes, the coverslips were mounted in DAKO (Agilent, S3023) on microscope slides and imaged on a Zeiss Axio Imager Z2. For microglial survival and cell type composition and analysis 20x magnification was used. For morphological and fluorescence analysis per cell x63 magnification with oil was used.

### Flow cytometry

Cultured cells were washed with 1xPBS and dissociated into a single-cell suspension using TrypLE (Gibco, 120604021). The cells were dissolved in Viability Dye eFluor 780 (Invitrogen, 65-0865-14,) and incubated at 4 °C for 30 min. Subsequently, cells were centrifugated at 300xg for 5 min and washed with flow buffer (1% BSA in PBS) twice. Then, cells were fixed for 15 min at RT with 2% paraformaldehyde followed by another wash. To prevent aspecific binding to Fc-receptors, the cells were blocked with 2.5% Human TruStain FcX blocking solution (BioLegend, 422302) for 10 min. Following antibodies were used: CD43-PE (1:100, BioLegend, 343203), CD45-FITC (1:200, eBioscience, 11-9459-42), CD45-PE/Cyanine7 (1:200, BioLegend, 304015), CD11b-PE (1:100, BioLegend, 301305), CX3CR1-APC (1:100, BioLegend, 34110), HLADR-APC (1:50, eBioscience, 47-9956-42), P2RY12-BV421 (1:100, BioLegend, 392105), CD206-APC (1:50, BD Biosciences, 550889), CD24-PE Cyanine7 (1:50, BioLegend, 311119), GFAP-BV421 (1:100, BioLegend, 644710).

After 30 min incubation at 4 °C, the cells were washed three times and finally dissolved in 200 µl flow buffer for flow cytometry measurement (Gallios, Beckman Coulter). Gating of side scatter (SS) area versus forward scatter (FS) area was used to define the cell population. Duplicates were removed by gating between FS height and FS Area. Further, dead cells were negatively gated by viability dye intensity. Unstained samples were used to determine the auto fluorescence levels of cells in the channels corresponding to the antibodies used. The compensation strategy was determined using beads (BioLegend, 424602). Analysis was performed using Kaluza C Analysis software (Beckman Coulter).

### RNA-sequencing

Plating and culturing procedure was done according to the specific cell type. Cultured cells were washed with 1xPBS and collected using DNA/RNA shield (Zymo Research; ZY-R1200-125). Human post-mortem microglia were stored in Trizol and received from the Netherlands brain bank. RNA was isolated using the Quick-RNA Microprep kit (Zymo Research, ZY-R1051) according to manufacturer’s instructions. After quality control, NEBNext® Ultra™ II RNA Library Prep Kit (Illumina) was used for library preparation as per manufacturer’s recommendations. Sequencing was performed on a NovaSeq X+ (Illumina) at 2x150bp configuration with coverage of 20 million reads per sample. Raw RNA-sequencing reads were processed using the nf-core/rnaseq pipeline (v3.18) implemented in Nextflow. Quality control was performed using FastQC and MultiQC. Adapter trimming was conducted with Trim Galore. Reads were aligned to the human reference genome (GRCh38) using STAR with default parameters. Transcript-levels were quantified with Salmon. Log2 transformation with a pseudo-count of +1 was performed on counts per million (CPM) to achieve normal distribution. Heatmaps were generated using Euclidean clustering in the *pheatmap* R package.

### Single-cell RNA-sequencing

Microglia monocultures and neuron-microglia co-cultures were differentiated in 6-well plates according to their protocol described above. The microglia monocultures also received PLO/human laminin coating as the co-cultures for fair comparison. The experiment was performed in technical triplicates per condition. At DIV 28 the cultures were dissociated using Accutase (20 min at 37 °C). All conditions were treated exactly the same to ensure a fair comparison, even though microglia monocultures did not require as long an incubation period as the very dense neuron-glia networks. The single-cell suspensions were processed using the Chromium Next GEM Single Cell Fixed RNA Sample Preparation Kit (10x Genomics, PN-1000414) according to the manufacturer’s guidelines. The samples were fixed for 24 hours at 4 °C and stored with Enhancer at -80 °C until further processing. Library preparation was performed using the Chromium Single Cell Flex Kit (10x Genomics) following the manufacturer’s protocol (CG000527 Rev F). All samples were incubated for 20 hours at 42 °C with an anti-human probe set. Each condition had a unique barcode to allow for multiplexing. After pooling the samples and subsequent washes, three lanes of a 10x Chip Q were loaded with each condition being represented in each final lane/library. Raw sequencing data were processed and de-multiplexed using the Cell Ranger pipeline (10x Genomics, Cellranger v. 9.0.0) with human reference genome refdata-gex-GRCh38-2024-A and Chromium_Human_Transcriptome_Probe_Set_v1.1.0_GRCh38-2024-A.

Downstream analyses, including quality control, normalization, library integration, doublet detection, clustering, and differential gene expression analysis, were performed using Seurat (v4.3.0)) in R. Quality control criteria included the removal of cells with fewer than 200 genes detected, fewer than 500 UMIs or with more than 10% mitochondrial gene content. To remove doublets, cells with a UMI count two standard deviations (above the mean were excluded. Further, doublets were identified by mixed gene expression profiles and their characteristic floating and small cluster shape. Harmony^48^ (v1.2.3) was used for integration of the three batches/libraries. Cell types were annotated based on the top differentially expressed genes of the Seurat clusters. The homeostatic gene sets and neuronal remodeling genes were derived from the current microglia nomenclature consensus^8^ and transcriptional phenotypes found in literature^22^. The environment-dependent signature (ex vivo induced) is based on the findings from Gosselin et al^16^. AD pathology and lipid processing related gene sets are derived from a recent study using post-mortem brain tissue about human microglial state dynamics in AD^38^. To predict ligand-receptor pairs based on the single cell transcriptomes and infer cell-cell communication CellChat (v2.1.2) was used^40^

### Micro-electrode array recording and analysis

Neuronal network activity was recorded using the Axion Maestro Pro system equipped with AxIS Navigator software (Axion Biosystems). On DIV0 hiPSCs were plated on CytoView 48-well micro-electrode arrays (MEAs) and then differentiation was initiated. To ensure stable temperature of 37 °C and 5% CO_2_ during measurements, all plates acclimatized for a minimum of 10 min before starting the recording. Recordings were always performed ∼24 hours after medium change as this procedure influences firing. Spikes were detected when voltage deflections exceeded an adaptive threshold of ±6 standard deviations from the noise level. Processing of the MEA data was done using the Axion Biosystems NeuralMetric Tool. Bursts were detected when at least 5 consecutive spikes had a maximum inter-spike interval of 100 ms each. The envelope network burst detection algorithm of was used with these settings: threshold = 1.5, minimum inter burst interval = 100 ms, minimum number of active electrodes = 70%, burst inclusion = 75%.

### Whole-cell patch clamp recordings

Coverslips were placed in a recording chamber on a microscope stage and were perfused with oxygenated (95% O_2_ /5% CO_2_) artificial cerebrospinal fluid (ACSF) at 32°C. ACSF contained (in mM): 24 NaCl, 1.25 NaH_2_PO_4_, 3 KCl, 26 NaHCO_3_, 11 glucose, 2 CaCl_2_ and 1 MgCl_2_. Pipettes for patch clamp recordings were pulled from borosilicate glass capillaries with filament and fire-polished ends (inner diameter 0.86 mm, outer diameter 1.05 mm, resistance 5–8 MΩ, Science Products GmbH) using a micropipette puller (PC-10, Narishige). For recording miniature excitatory postsynaptic currents (mEPSCs) and AMPA receptor-mediated currents, pipettes were filled with a filtered intracellular solution containing (in mM): 30 potassium gluconate, 5 KCl, 10 HEPES, 2.5 MgCl_2_, 4 Na_2_-ATP, 0.4 Na_2_-GTP, 10 Na_2_-phosphocreatine and 0.6 EGTA. The pH was adjusted to 7.2. Osmolarity was adjusted to 290 mOsmol using a Semi-Micro Osmometer (K-7400, Knauer). Recordings were carried out in voltage-clamp mode, acquired using a digitizer (1140A, Digidata) and an amplifier (Multiclamp, 700B, Molecular Devices), with a sampling rate at 20 kHz and a lowpass filter at 1 kHz. Recordings were not corrected for liquid junction potential (−15.646 mV). Recordings were not analyzed further if series resistance was >25 MΩ or when it became >10% of the membrane resistance. mEPSCs were recorded for 10 min in the presence of 1 µM tetrodotoxin (Tocris, 1069) and 100 µM picrotoxin (Tocris, 1128) at a holding potential of -60 mV at DIV 49. To measure light-evoked synaptic currents, we sparsely transduced neurons with a lentiviral vector containing channelrhodopsin-2 (ChR2) tagged with an enhanced yellow fluorescent protein (eYFP) under a human Synapsin I promoter (Addgene, 20945) at DIV 1. At DIV 28, before patching a neuron, an illumination of 470 nm LED light (KSL2, Rapp OptoElectronic) was given to verify whether the neuron was positive or negative for ChR2. A ChR2-negative cell was recorded in voltage clamp mode at a holding potential of -60 mV and +40 mV. Light-evoked synaptic responses were induced by exciting nearby ChR2-positive cells using a 5 ms illumination with 470 nm LED light through the objective lens (60x). For the final traces, 30 sweeps were averaged. Light-evoked synaptic responses were 2,3-Dioxo-6-nitro-1,2,3,4-tetrahydrobenzo[*f*]quinoxaline-7-sulfonamide (NBQX)-sensitive (50 µM; dissolved in DMSO; Tocris, 0373) indicating these currents were AMPA receptor-mediated (Suppl. Fig. 4). In case a ChR2-positive cell was recorded, the location of the stimulus was adjusted to avoid excitation of the patched neurons.

## Acknowledgments

We thank all members of the de Witte and Nadif Kasri labs, as well as Marijn Kuijpers for their valuable input during meetings and discussions. We thank Jorge Domínguez Andrés for kindly contributing the human monocyte samples. We also thank the RadboudUMC Flow Core for supporting experiments as well as providing broncho-alveolar fluids. This work was supported by the Hypatia Grant from the RadboudUMC, the Simons Foundation (00010410) and the Netherlands organization for health research (ZonMW Open Competitie 09120012110034).

---

**Supplementary figure 1.**
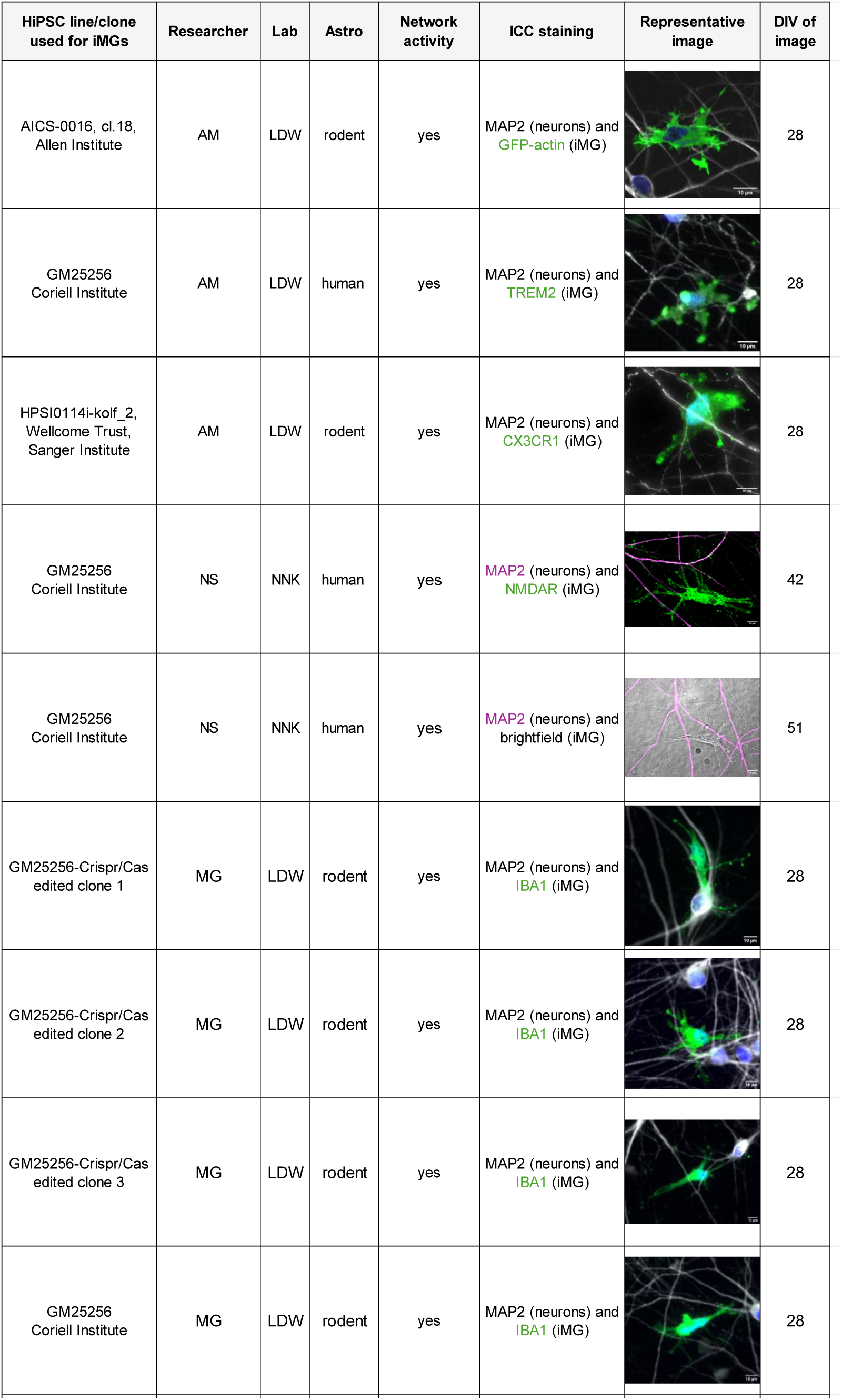
HiPSC lines tested for iMG differentiation in co-maturation protocol by different researchers. Astro = Astrocytes, ICC = Immunocytochemistry, scale bars = 10 μm.

**Supplementary figure 2.**
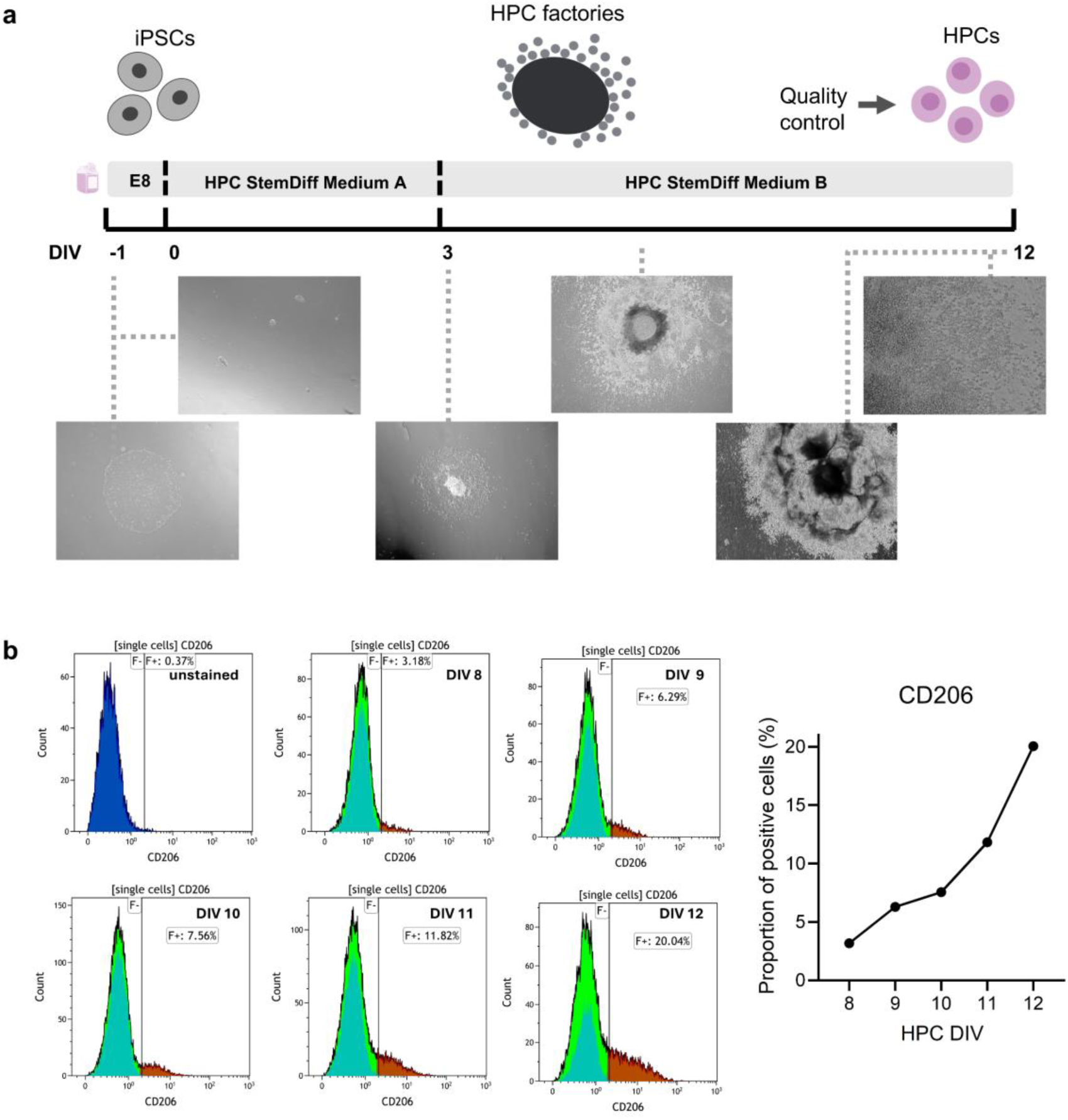
Timeline of HPC differentiation. **(a)** Schematic protocol overview for generation of HPCs from hiPSCs with representative brightfield images of the cultures. **(b)** Flow cytometry analysis of CD206 expression in HPC population sampled at DIV 8 to DIV 12 along the differentiation timeline, N = 1.

**Supplementary figure 3.**
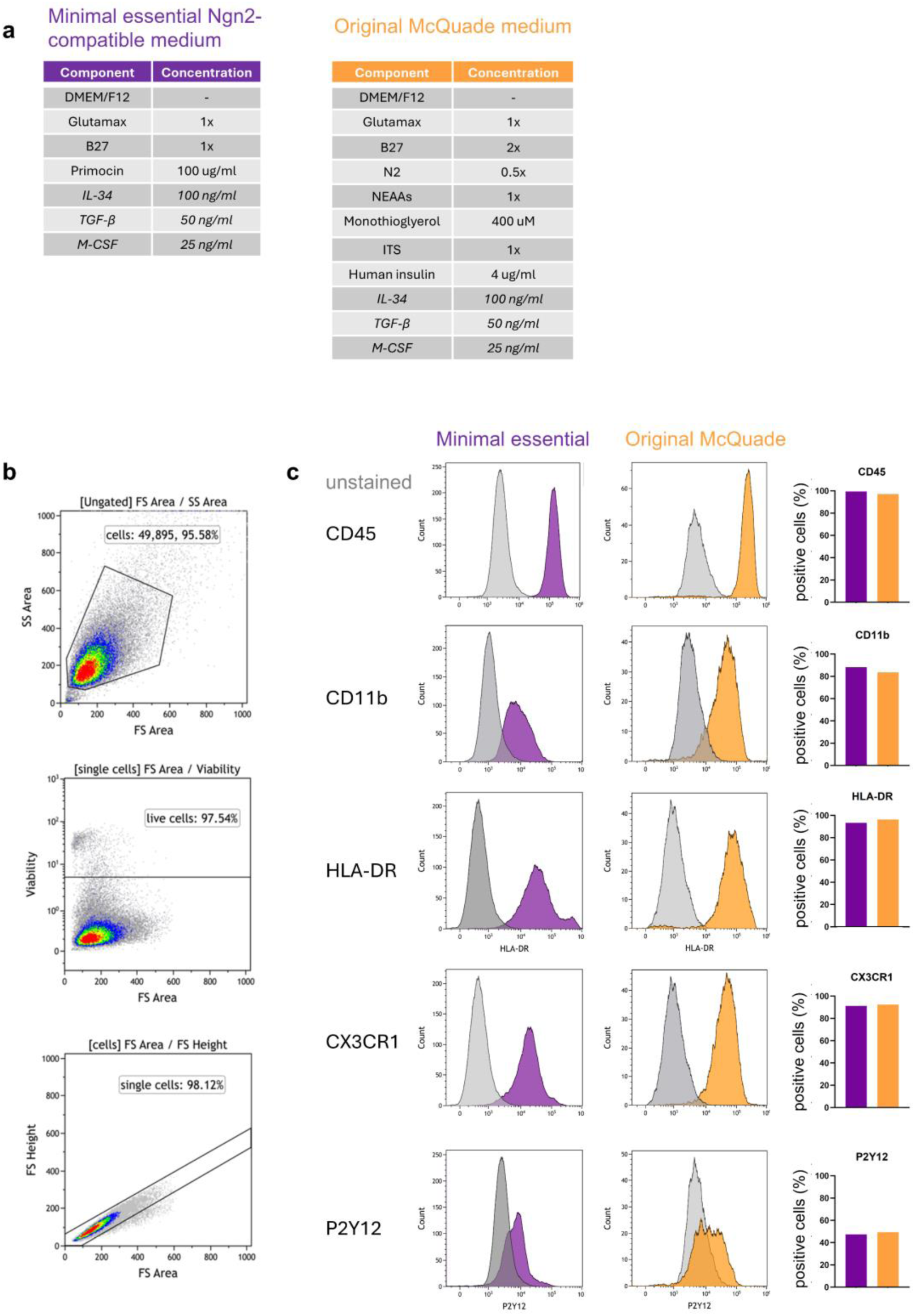
Comparison original McQuade medium to minimal essential medium. **(a)** Detailed medium composition of the new minimal essential medium compatible with *Ngn2-*induced neurons and the original medium based on McQuade et al. (2018). **(b)** Representative Flow Cytometry gating of iMG cultures. **(c)** Flow cytometry analysis of CD45, CD11b, HLA-DR, CXC3CR1, and P2Y12 in iMGs differentiated with the two different medium compositions, N = 1.

**Supplementary figure 4.**
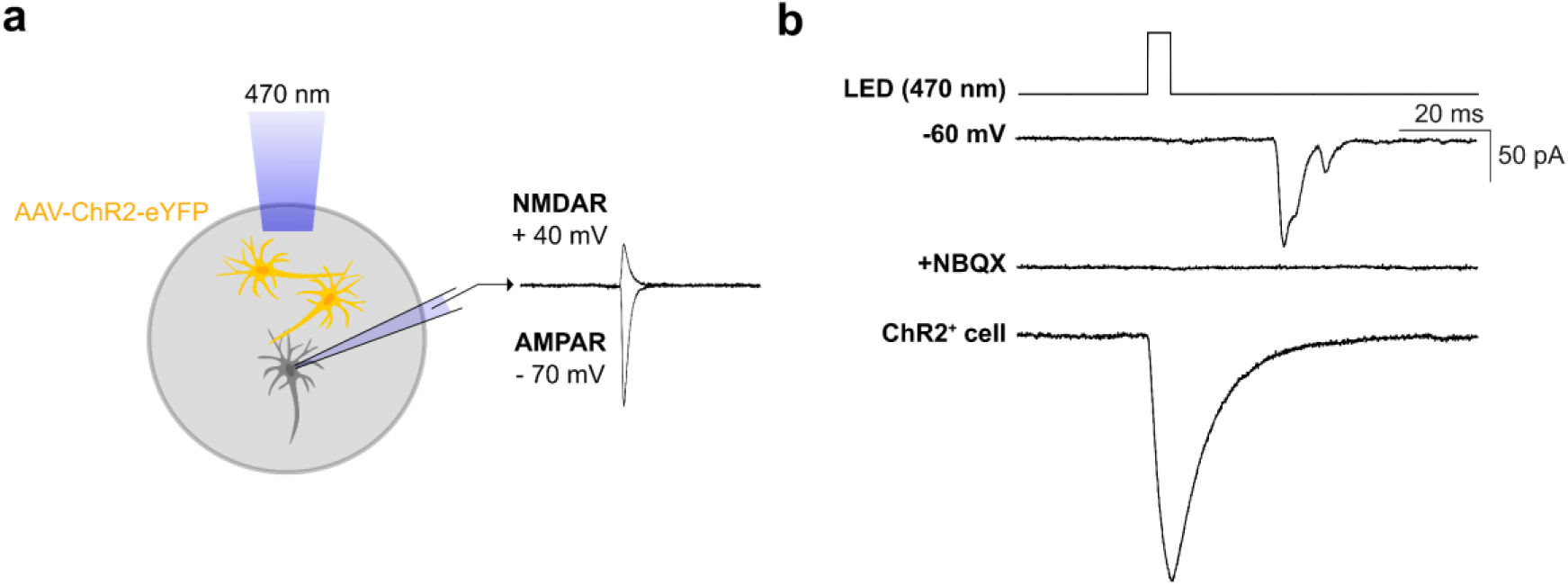
Patch clamp experiment for optogenetically induced synaptic responses. **(a)** Schematic overview of experimental design for optogenetics to induces light-evoked synaptic responses using Channelrhodopsin (ChR2). **(b)** Representative traces of optogenetically induced synaptic response showing no response when NBQX was washed in.

**Supplementary figure 5.**
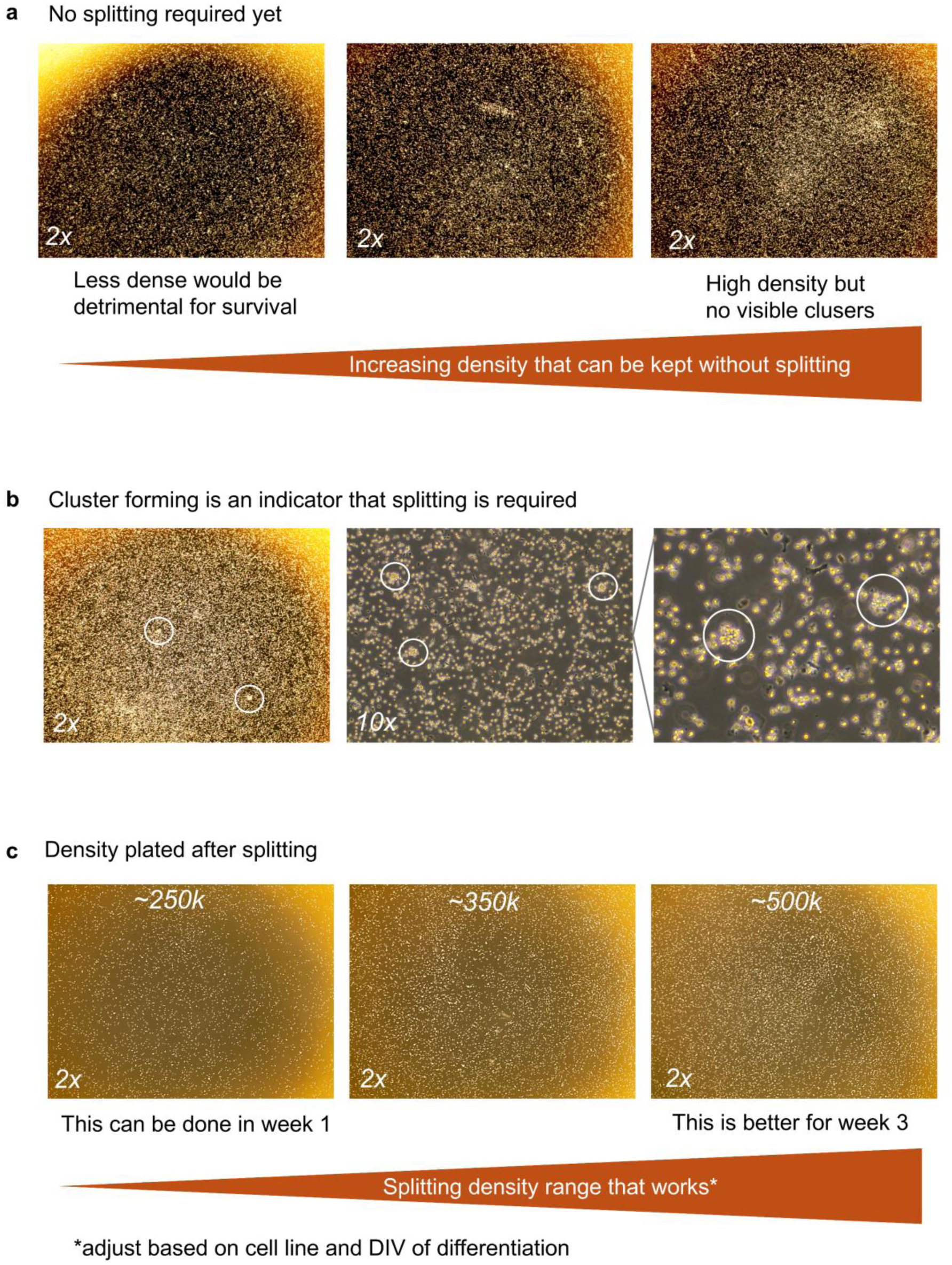
Recommendations on splitting iMGs. Representative brightfield microscope images (2x magnification) of iMGs **(a)** that do not require splitting yet, **(b)** forming clusters as an indicator that splitting is required and **(c)** after splitting.

